# Benchmarking long-read variant calling in diploid and polyploid genomes: insights from human and plants

**DOI:** 10.1101/2025.05.14.653922

**Authors:** Yoshinori Fukasawa

## Abstract

Accurate characterization of genetic variation is fundamental to genomics. While long-read sequencing technologies promise to resolve complex genomic regions and improve variant detection, their application in complex genomes has not been well validated. Here, we systematically investigate the factors influencing variant calling accuracy using accurate long reads. Using human trio data with known variants to simulate variable ploidy levels (diploid, tetraploid, hexaploid), we demonstrate that while variant sites can often be identified accurately, genotyping accuracy decreases with increasing ploidy due to allelic dosage uncertainty. This highlights a specific challenge in assigning correct allele counts in polyploids even with high depth, separate from the initial variant discovery. We then assessed genotyping and variant detection performance in real genomes with varying complexity: the relatively simple diploid *Fragaria vesca*, the tetraploid *Solanum tuberosum*, and the highly repetitive diploid *Zea mays*. Our results reveal that overall variant calling accuracy is influenced strongly by inherent genome complexity (e.g., repeat content).

Furthermore, we identify a critical mechanism impacting variant discovery: structural variations between the reference and sample genomes, particularly those containing repetitive elements, can induce spurious read mapping. This effect is likely exacerbated by the length and accuracy of long reads. This leads to false variant calls, constituting a distinct and more dominant source of error than allelic-dosage uncertainty. Our findings underscore the multifaceted challenges in long-read variant analysis and highlight the need for ploidy-aware genotypers and bias-aware mapping strategies to fully realize the potential of long reads in diverse organisms.

## Background

The comprehensive identification and genotyping of genetic variants, including single nucleotide variants (SNVs), small insertions/deletions (indels), and structural variations (SVs), are crucial for understanding genotype-phenotype relationships, population genetics, and evolutionary processes [1]. The advent of long-read sequencing technologies, such as Pacific Biosciences (PacBio) and Oxford Nanopore Technologies (ONT), has significantly advanced variant detection capabilities. Long reads can span repetitive regions, resolve complex structural rearrangements, and facilitate haplotype phasing, overcoming limitations inherent to short-read sequencing [2, 3]. Recent advances in third-generation long-read sequencing (e.g., PacBio HiFi and ONT duplex reads) have enabled accurate mapping across repetitive and structurally complex regions, improving variant calling performance in previously challenging genomic contexts [4, 5].

Despite these advantages, accurate variant calling in non-model organisms, particularly those with polyploid or highly complex genomes, remains a significant hurdle [6].

Polyploidy, the state of having more than two complete sets of chromosomes, is widespread in plants and also occurs in some animals and fungi. It introduces challenges related to discriminating between homologous, paralogous, and homeologous sequences, and accurately determining allelic dosage (the number of copies of each allele at a heterozygous site) [7]. Although recent studies have benchmarked variant calling performance using long-read data, these efforts have largely focused on diploid genomes or structural variants [8, 9]. Cooke et al. previously evaluated short-read variant callers on synthetic polyploid datasets [10], providing valuable insights into genotyping challenges under polyploidy. However, systematic benchmarking of small variant calling using high-accuracy long-read technologies in such contexts remains largely unexplored.

Genome complexity, encompassing factors like high repeat content, elevated heterozygosity, and large repertoires of SVs relative to the reference, further complicates variant analysis irrespective of ploidy [11]. These features can lead to ambiguous read mapping, collapsed assemblies, and erroneous variant calls [12]. While long reads are expected to ameliorate some mapping issues associated with repeats, the interplay between ploidy, inherent genome complexity, and the accuracy of long-read based variant calling requires systematic investigation. Specifically, it is not yet clear whether ploidy or genome complexity plays a more significant role in determining performance in the context of long read.

Furthermore, reference genomes often represent only one haplotype or accession, while significant structural variation exists between individuals or accessions within a species [13]. To address this, the construction of pangenomes is actively being pursued. However, for many species where high-quality pangenomes are not yet available, the use of a single reference genome remains a practical necessity. Such divergence, particularly involving structural variations and repetitive elements, can compromise long-read mapping accuracy and downstream variant detection. A better understanding of these effects is therefore essential.

Variant calling in polyploids is notably complicated by issues such as read mapping errors and allelic dosage uncertainty stemming from their complex genomic nature [14]. However, despite the promise of long-read sequencing to potentially mitigate some of these challenges, the specific impact of polyploidy on long-read variant calling accuracy remains largely unexplored.

Accurate long-read sequencing has reached a mature stage for diploid human genomes, with its high precision making it a promising tool for clinical applications [15]. While the significance of this progress is evident in the medical context [16], broader applications in genomics demand robust performance in more complex settings such as higher-order polyploid organisms or species with high levels of intraspecific variation. Yet, the reliability of variant calls in such contexts and the contributing sources of error have yet to be systematically characterized.

To address this, we investigated genotyping performance across different levels of ploidy and genome complexity. We first evaluated challenges associated with polyploidy using synthetic human data, then extended our analysis to real datasets from three plant species with varying genomic complexity. Finally, we assessed how reference-sample differences influence downstream variant calling performance.

## Results

### Benchmark Design and Read-alignment performance

In this study, we included plant genomes alongside the human genome to evaluate performance across species with high-quality reference assemblies but differing in genome size and sequence complexity. *Fragaria vesca* (*F. vesca*) was selected as a model of a relatively simple genome: it is diploid, has a small genome size and low repeat content, and exhibits low heterozygosity due to its selfing nature. In contrast, *Zea mays* (*Z. mays*) was chosen as a representative of large, highly repetitive genomes. Despite its complexity, it has established inbred lines, which minimize heterozygosity. Lastly, we selected *Solanum tuberosum* (*S. tuberosum*) as a polyploid model. Its genome has moderate repeat content and is characterized by high haplotype diversity, providing a useful contrast in terms of ploidy and allelic variation.

Before turning to plants, we established a baseline using synthetic polyploids derived from human genomes. By confirming that variant-detection performance remains robust in this controlled setting, we could later attribute any additional error patterns in plants to sequence features. For all three species, high-accuracy long-read sequencing data derived from individuals genetically distinct from the reference were publicly available and therefore used in this study (Table S1). The same aligner and parameter set (see Methods) were applied across all datasets, ensuring that downstream comparisons were not confounded by disparate mapping conditions.

To ensure downstream comparisons rested on solid foundations, we performed a uniform validation of the read alignments. All datasets exceeded standard benchmarks for long-read WGS: overall mapping rates were > 98 %. The distribution of mapping quality followed the expected complexity gradient, highest in Human and *F. vesca*, intermediate in *S. tuberosum*, and lower in *Z. mays*, providing adequate positional reliability for downstream variant calling (Table S2).

### Genotyping Ambiguity in Polyploids

To distinguish between sequence-based features and allele-dosage uncertainty arising from polyploidy, we performed evaluations using the well-established human trio genome as a standard reference. Following the approach established in a previous study [10], we constructed synthetic polyploid test samples with known truth using the Ashkenazi trio (HG002/HG003/HG004 from Genome in a Bottle) (see Methods). In brief, we merged the parental genomes to simulate an autotetraploid (4x) and combined both parents plus child for a hexaploid (6x) genomes. PacBio HiFi reads for the relevant individuals were pooled and randomly subsampled to generate 10, 30, 50, 70, and 90× coverage for each synthetic polyploid. Variants were then called on these datasets assuming ploidy 2, 4, or 6. A comprehensive summary of performance metrics across all species and variant callers is provided in Supplementary Table S3.

Although performance declines as ploidy increases, ensuring sufficient coverage per haploid genome can partially mitigate this loss (Figure 1). GATK performance was comparable to, and only marginally better than, the Illumina results reported in prior studies [10]. Overall, GATK maintained high genotyping accuracy in diploids and, given adequate coverage, showed only a modest decline even in tetraploids. For FreeBayes, performance in diploids showed little variation, but it deteriorated markedly as ploidy increased (Figure 2). Although sufficient sequencing depth is important, the results suggest additional factors that coverage alone cannot compensate for. Variant-type stratification revealed that FreeBayes’s performance decline is driven largely by indels (Figure S1 and S2). Indels are more susceptible than SNVs to platform-specific sequencing biases [2], which likely accounts for the reduction.

**Figure 1.**
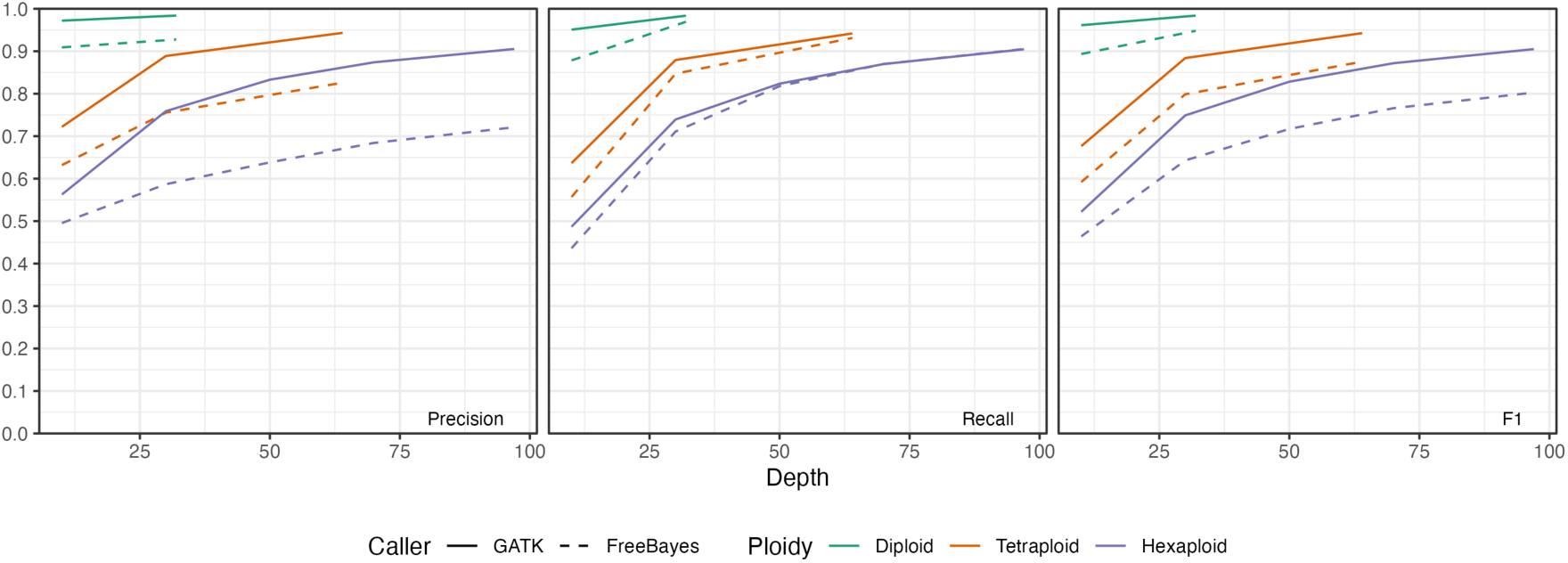
Performance of small variant genotyping on the synthetic human polyploid genome using high-accuracy long reads. Precision, recall, and F1 scores are presented.

**Figure 2.**
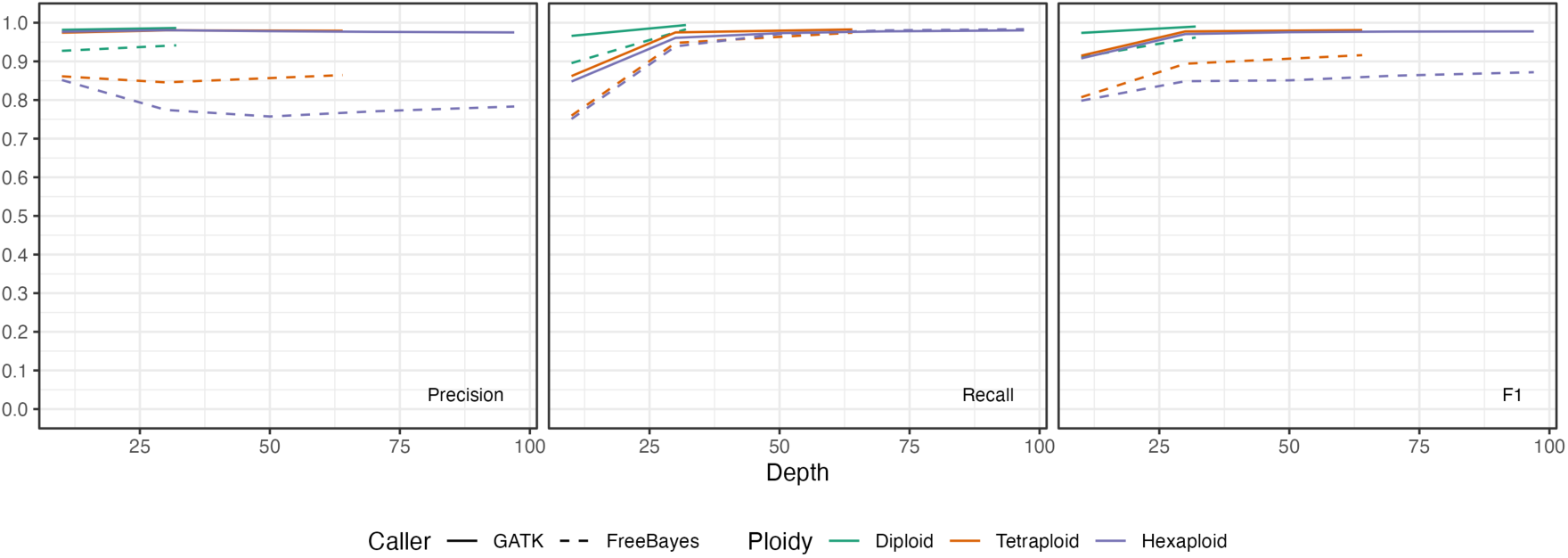
Performance of small variant detection on the synthetic human polyploid genome using high-accuracy long reads. Precision, recall, and F1 scores are presented.

Allele detection exhibited the same pattern. With GATK, reliable detection was achievable at lower sequencing depths, and once a moderate depth was reached, accuracy remained high irrespective of ploidy level (Figure 2). For allele detection, performance for both SNVs and indels reaches a plateau at about 30 × coverage (Figure S3). In tetraploids, however, raising coverage beyond this level for indels correlates with the calling of spurious variants and, intriguingly, a slight decline in precision (Figure S4).

Critically, one of major errors in the polyploid calls was genotyping errors. The variant was often flagged by the caller, but the exact genotype (allelic copy number) was mis-assigned. This aligns with theoretical expectations – as ploidy rises, each haplotype is covered by fewer reads on average, making it harder to distinguish a real low-frequency variant from sequencing error noise [14]. Nonetheless, the fact that precision and recall only dipped a few percentage points by tetraploidy indicates that allelic dosage uncertainty, while present, is manageable with GATK at typical coverage. A further salient issue is the difficulty of genotyping indels, a limitation most pronounced with FreeBayes. By contrast, for SNVs FreeBayes performs on par with, or marginally better than, GATK. SNV detection was reliable with both tools, and GATK proved more robust for indel calls. Encouraged by evidence that performance remains strong even in highly polyploid genomes, we next constructed high-confidence truth sets for three plant genomes that span a broad range of genomic complexity.

### Extending Variant Calling Benchmarks to Diverse Plant Genomes

By employing synthetic polyploids derived from human genomes, we confirmed that variant detection performance remains robust and established a baseline for evaluating genotyping accuracy across increasing ploidy levels. Using these results as a reference, we further evaluated performance on genomes with sequence characteristics distinct from those of humans: *F. vesca* (220 Mb, diploid), *S. tuberosum* (840 Mb, tetraploid), and *Z. mays* (2.1 Gb, diploid but highly repetitive).

To define high-confidence variants relative to the reference genome, we obtained high-quality genome assembly based on long-read sequencing data (Table 1). The same long-read data, derived from the individuals used for assembly, were also used for read mapping (Table S1) and variant calling throughout this study. Although completely eliminating sequencing errors is still challenging, recent genome assemblies have expected error rates below 1%, a level sufficiently low that it does not substantially impact the performance evaluations conducted in this study.

**Table 1.** Summary of plant genomes used for benchmarking analyses.

| Species<br>(common name) | Ploidy | Assembly version<br>(accession/Database) | Haploid size (Mb) | Estimated<br>repeat<br>content (%) | Reference |
| --- | --- | --- | --- | --- | --- |
| *Fragaria vesca*<br>(woodland<br>strawberry) | 2 × | drFraVesc1<br>(GCA_964146915.1/G<br>CA_964165485.1) | 220 | ~35 | [22] |
| *Solanum tuberosum*<br>(potato) | 4 × | Otava v1.0<br>(Spud DB) | 840 | ~66 | [21, 23] |
| *Zea mays*<br>(maize) | 2 × | Mo17 CAU T2T<br>(GCA_022117705.1) | 2,100 | ~88 | [20] |

We performed chromosome-level comparisons between these highly accurate genome assemblies and representative reference sequences from the same species. A similar approach has already been adopted for benchmarking human genomes [17, 18]; with the recent availability of high-quality genome assemblies, this methodology has become increasingly practical for wide range of organisms. By delineating one-to-one alignable regions we created a mask that limits all downstream evaluations to parts of the genome where both alleles are confidently resolved.

Within the high-confidence regions we established variants as benchmarks for subsequent analyses (Figure 3A-C). We further strengthened this benchmark by evaluating variants using Merfin’s k-mer-based approach. An independent assessment confirms that more than 99% of variants are supported by *k*-mers (Table S4), which validates both the quality of the assembly used in this study and the variants detected from it, independently of alignments.

**Figure 3.**
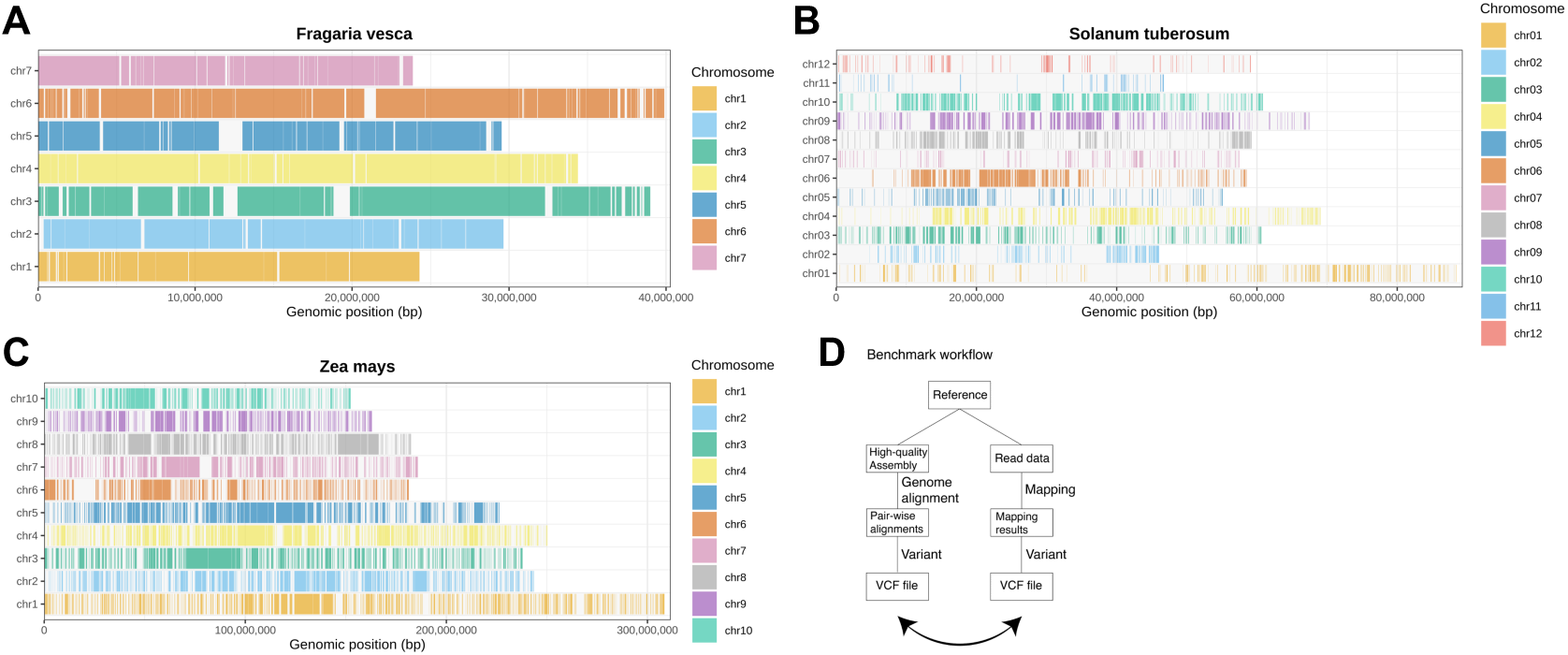
Schematic diagram of confidence regions defined by assemblies from individuals of the same species relative to the reference genome. For each chromosome, confidence regions that are uniquely aligned to the reference and consistently represented across all haplotypes are shown as colored rectangles. (A) *Fragaria vesca*, (B) *Solanum tuberosum*, (C) *Zea mays*, (D) Schematic overview of a semi-automated method for benchmark data.

*F. vesca* is a small genome species with low reported intraspecific diversity [19], and our analysis confirmed a high degree of similarity between the reference genome and the sequenced sample (Figure 3A). In contrast, *Z. mays* is known for its high intraspecific diversity, and the proportion of one-to-one alignable regions between the reference and sample genomes was intermediate (Figure 3B). This is consistent with previous studies [20]. For *S. tuberosum*, the observed variation is largely attributable to the diversity among its four homeologous chromosomes (Figure 3C). Our findings agree with earlier reports showing limited core regions shared across all haplotypes [21].

### Variant-calling Accuracy Across Plant Species

To test performance in real-world diverse genomes, we applied the long-read variant calling pipeline to the three plant genomes of increasing different complexity (Figure 3D). We aligned reads to the respective reference genomes (the *Fragaria vesca* v6.0, the *Solanum tuberosum* DM1-3 516 R44 v6.1, and the *Zea mays* B73 v5 reference genome). Variant calling was performed with both GATK and FreeBayes (with ploidy set to 2 for strawberry and maize, 4 for potato). We evaluated the resulting variant calls against a set of putative truth variants for each species (see Methods for truth set derivation).

Genotyping performance considerably differed across genomes, reflecting species-specific genomic properties. For the simplest genome, *F. vesca*, we obtained accuracy on par with human benchmarks: Overall F1 was comparable to that observed in the human dataset, with SNV F1 reaching roughly 95 % (Figure 4, S5–S7). This suggests that a high-quality diploid plant genome of small size can be surveyed for variants with nearly the same confidence as a human genome using long reads. The errors that did occur in *F. vesca* were mainly in small repetitive sequences and collapsed tandem repeats. In tetraploid potato, performance was clearly reduced relative to *F. vesca* and synthetic-tetraploid human results (Figure 4, S8-S10). In addition to being an autotetraploid, the potato genome exhibits substantial genomic divergence between the Otava and DM genotypes, likely explaining the notably reduced recall in variant detection. Finally, maize showed the greatest challenges, even though the comparison was between two diploid inbred lines.

**Figure 4.**
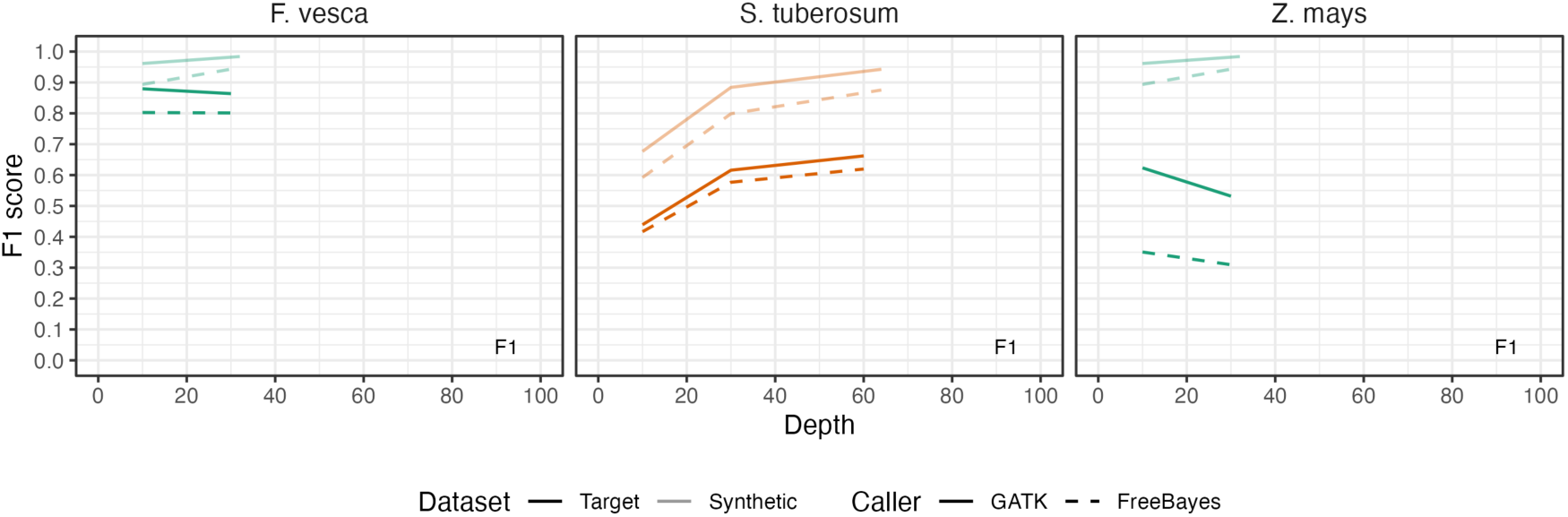
Performance of small variant genotyping (F1 score) in *Fragaria vesca*, *Solanum tuberosum*, and *Zea mays*. The corresponding results for the synthetic human datasets are displayed as a semi-transparent reference trace. F1 scores for each species are presented.

Despite the absence of polyploidy or high heterozygosity, its extreme repeat content and structural diversity led to the most substantial performance drop. Genotyping performance in *Z. mays* fell sharply compared with diploid human and *F. vesca (*Figure 4, S11-S13). In fact, the performance was even lower than that observed for potato, an autotetraploid species with high sequence divergence, suggesting that factors other than ploidy may play a critical role in genotyping accuracy. The decline was driven mainly by lower precision— that is, a large increase in false-positive calls (Figure S11-S13). It exhibited a pronounced reduction even in SNV precision, falling to below 50 % (Figure S12). Under the same analysis pipeline, *Z. mays* produced false-positive calls at rates tens of times higher than those observed for either the human or *F. vesca* genomes. The trend was clear – as genome size, repeat content, and structural heterogeneity increased, variant calling accuracy decreased. Because *F. vesca* and inbred lines of *Z. mays* are highly homozygous diploid genomes, the likelihood of genotyping errors was expected to be low. Consequently, we focused on whether each variant was correctly detected rather than on evaluating genotype accuracy.

The observed trend became more pronounced: while recall remained high as anticipated, limited precision emerged as a consistent issue across species (Figure 5, S14-S22). This limitation was particularly evident in *Z. mays*. For SNVs, recall closely matched values of human genome benchmarks, indicating potential practical applicability (Figure S15, S18, S21). However, precision for even SNVs remained substantially lower in *S. tuberosum* and *Z. mays* (Figure S15, S18, S21). The high recall and precision in variant detection observed for *F. vesca*, in contrast to the markedly reduced precision seen in *Z. mays*, highlight notable differences in variant calling performance across species. These findings suggest that the observed discrepancies are primarily driven by genome-specific sequence characteristics rather than limitations of the variant detection pipeline or errors due to solely polyploidy.

**Figure 5.**
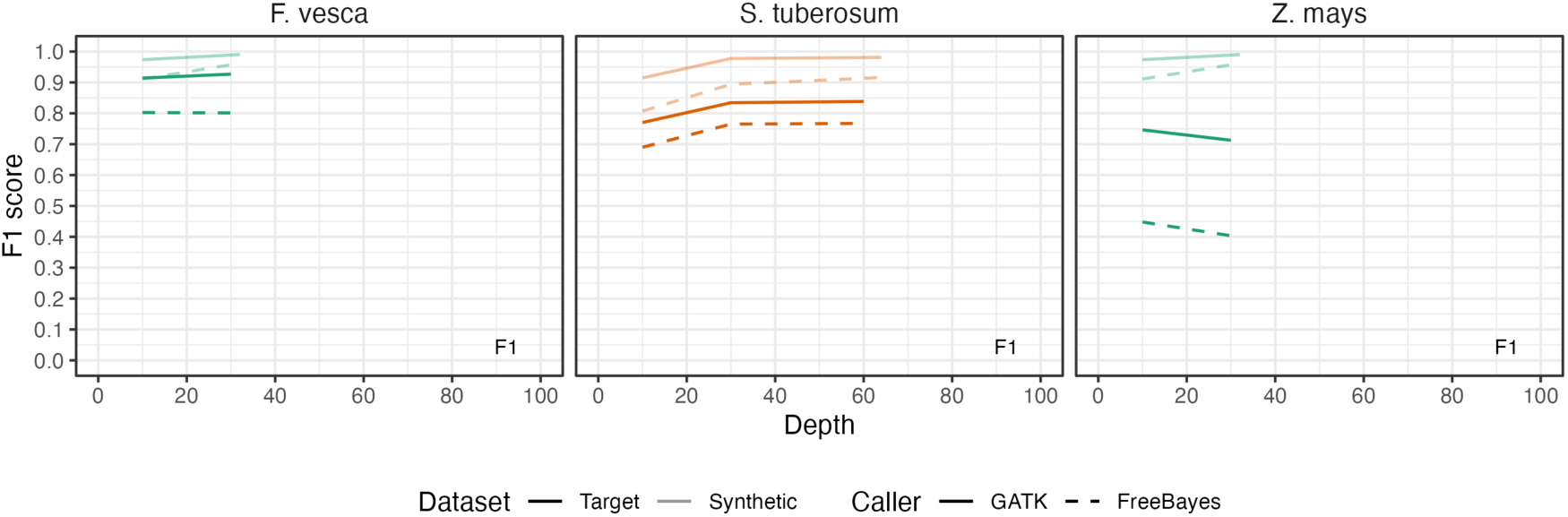
Performance of small variant detection (F1 score) in *Fragaria vesca*, *Solanum tuberosum*, and *Zea mays*. The corresponding results for the synthetic human datasets are displayed as a semi-transparent reference trace. F1 scores for each species are presented.

Notably, the *maize* results underscore that a highly repetitive, structurally variable diploid can be even more challenging for variant calling than a less repetitive polyploid even using accurate long reads. The maize line Mo17 (an inbred distinct from B73) carries many structural variants relative to the B73 reference [20]. Decades of maize genomic research have documented that any two maize lines can differ by presence/absence of thousands of genomic segments [24]. Our variant calling analysis reflected this: many false positives in maize were in regions where the sample’s reads were erroneously aligned due to sequence present in Mo17 but absent in the B73. In potato, the reference genome is a doubled monoploid line (essentially a haploid-derived assembly) that is divergent from the heterozygous tetraploid cultivar Otava [21]. Therefore, potato also exhibited reference-mismatch issues, though on a smaller scale than maize. In summary, our analyses of plant genomes demonstrate that, with residual read and base calling errors largely mitigated by accurate long read sequencing (i.e. HiFi), the principal constraint on variant-calling accuracy is how faithfully the reference genome reflects the sample genome’s repetitive and complex architecture.

### Reference bias and absent sequences are the dominant sources of errors

Even with long reads, alignment issues caused by an imperfect reference genome emerged as a primary contributor to false variant calls and missed variants. In the human dataset, which uses a high-quality reference (GRCh38) and where the sample is relatively well represented by that reference, difficult regions were limited. In the case of non-human organisms, a more significant concern likely arises from biases caused by differences between the reference genome and the genome under investigation. Whenever the sample genome contained an insertion or deletion not present in the reference, the aligner could introduce misalignments, causing reads to become gapped or clipped [14]. These misalignments can result in incorrect variant calls, not due to true sequence variation but as artifacts introduced during alignment. This reflects a general form of reference bias, in which reads are preferentially aligned to the reference structure even when it diverges from the sample genome [25]. Given concerns that such reference-bias may pose practical challenges in non-human genomes [14], we proceeded with further validation analyses using the selected genomes.

To better understand the errors in the plant genome variant calls, we performed a detailed error analysis, focusing on false positives unique to the long-read calls. A consistent finding was that many false SNV/indel calls were clustered rather than randomly distributed. In the maize dataset, for example, we observed clusters of false SNVs in the vicinity of large presence/absence variants. One illustrative case was a ∼55 kb SV present in the chromosome 1 of Mo17 but absent in B73 reference (Figure 6). In the alignment, reads from the large variant site partially aligned into the reference sequence. The variant caller then reported numerous SNVs and small indels in that region, all of which were artifacts resulting from reads originating from sample-specific regions that are absent in the reference genome. Essentially, the alignment constrained the reads to fit the reference, and the variant caller interpreted the resulting mismatches as true variants. This is a classic case of reference bias leading to false positives. Such alignment artefacts were a leading cause of false variant calls in maize and also contributed in the potato analysis. Although far less common, clusters of these false-positive SNVs were also detected in *F. vesca*.

**Figure 6.**
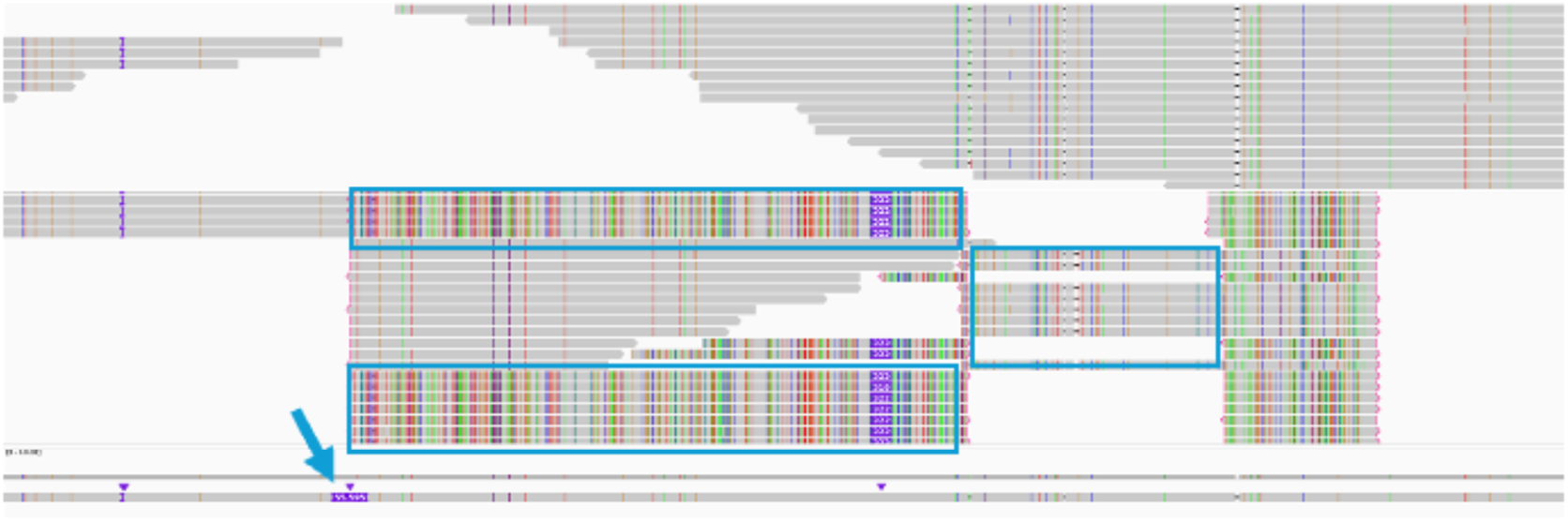
Repeat sequences originating from a large structural variant evade existing filters and become mixed with true alignments. The light-blue arrow marks a 55 kbp insertion in Mo17, and the light-blue box highlights reads derived from this insert.

In some regions, we observed clusters of apparent SNVs and indels that were ultimately determined to be artifacts. While we initially expected such regions to be associated with low mapping quality, a notable fraction of the supporting reads still carried high MQ scores (e.g., MQ = 60). This suggests that the aligner considered these placements to be reasonably confident, even though the underlying sequence was not present in the reference genome. These mismappings are likely exacerbated by the highly repetitive nature of plant genomes. Because maize is ∼88% transposable elements [26], many of these structural differences occur in repetitive contexts. If a repeat element is present in the sample but absent at the corresponding locus in the reference, reads may align to a homologous repeat elsewhere or in an incorrect orientation, leading to spurious variant calls.

To verify this directly, we simulated reads from the Mo17 genome and subjected them to the same analytical workflow. We produced whole-genome data at ∼10 and 30× coverage whose length and quality profiles matched those of PacBio HiFi reads. A read was retained if more than 50 % of its bases derived from reliable regions, defined as genomic intervals showing clear one-to-one sequence homology between Mo17 and B73. These reliable regions were identified through whole-genome alignment and were used to minimize reference bias by restricting analyses to sites with unambiguous correspondence between the reference and query genomes. Reads that failed to meet this threshold were discarded, as they may originate from presence/absence variation (PAV) segments. Because this threshold is deliberately conservative, the recall is lower than would be expected for a diploid genome; the critical point, however, is that the precision rises to the desired level (Figure 7). Furthermore, because alignments that would otherwise generate false positives have been filtered out, the atypical decline in performance observed at high read depth (30x) has been also eliminated.

**Figure 7.**
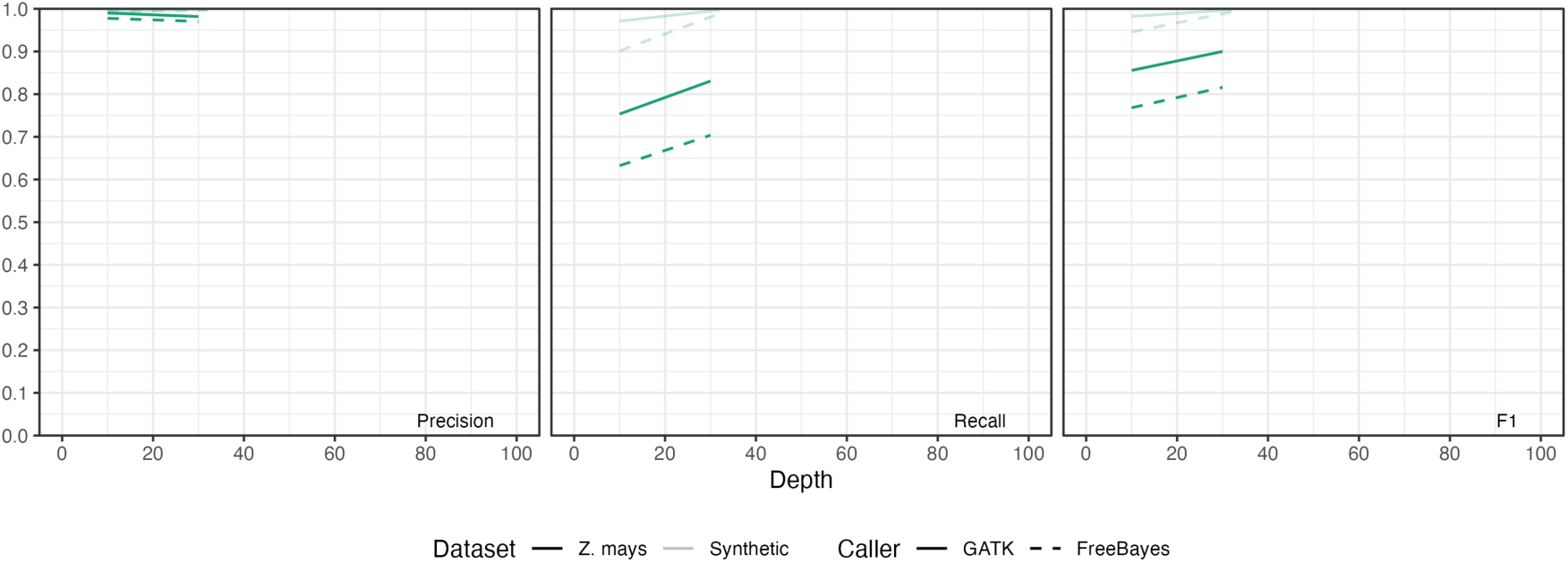
Simulated reads originating from reliable regions achieved the expected precision and improved recall for SNV genotyping. Precision, recall, and F1 scores are presented.

Our analyses indicate that sequence divergence between the reference and sample, especially in repetitive regions, is a major driver of false variant calls. This finding underscores the importance of high-quality reference genomes that capture population diversity. In crops like maize, where each accession can have unique sequence content not in the reference (and vice versa) [24], a single reference genome will inevitably cause reference-bias artifacts in variant calling. Long-read sequencing improves read placement compared to short reads, but as our results show, it does not fully resolve the reference divergence problem. Addressing these issues will require improved variant-calling algorithms and more flexible reference representations to minimize spurious calls that could mislead downstream analyses.

Taken together, these results suggest that most errors stem not from polyploidy per se, but from species-specific genome features such as large-scale presence/absence variation and high repeat content.

### Practical recommendations and limitations

It is important to note that DeepVariant was excluded from our main benchmark because it does not natively support genotyping in polyploid genomes. Although DeepVariant has demonstrated state-of-the-art performance in diploid human datasets, particularly for SNV and indel calling [27], its current implementation is not designed to accommodate more than two alleles per locus.

Nonetheless, given its strong performance in diploid systems, we conducted a pilot evaluation of DeepVariant on a tetraploid Solanum tuberosum dataset using its default diploid model. Surprisingly, the overall F1 score was slightly lower than that of GATK (Table S5), despite DeepVariant’s typically high precision. The drop in recall was primarily due to missed variants in genotypes with a single-copy alternative allele (e.g., AAAa), where approximately 60% of such sites were misclassified as homozygous reference (Table S6) at 60x depth. This likely reflects the fact that the model was trained exclusively on diploid data, leading to the misinterpretation of low-frequency alternative alleles as sequencing noise.

These observations underscore that extending deep learning-based variant callers to polyploid genomes will require not only revised genotyping logic but also retraining on polyploid-specific datasets. While such modifications are nontrivial, they represent a promising direction for future development, and we consider DeepVariant an important candidate for continued exploration in this space.

While long-read variant calling offers substantial advantages in genome-wide resolution, its computational cost remains nontrivial (Table S7). This cost increases approximately linearly with read depth, making high-coverage datasets particularly demanding in terms of runtime. Moreover, we observed a marked decline in indel-calling performance in all genomes. In such cases, we recommend using GATK for SNV calling, which demonstrated consistent and better performance even in complex polyploid contexts.

## Discussion

Our study demonstrates both the power and the remaining limitations of long-read sequencing for small variant detection in complex genomes. Notably, we show that PacBio HiFi enable highly accurate SNV calls even in polyploid genomes. In our human-based simulations, variant detection performance remained accurate from diploid to hexaploid, indicating that variant calling algorithms can accommodate the additional allele diversity with minimal loss of recall or precision. This is an encouraging result for researchers working with polyploid species, as it suggests that the fundamental task of identifying SNVs is not inherently compromised by increased ploidy. We found that genotype (allelic dosage) errors are the main issue in polyploid calling, rather than outright missed variants. This means that for many applications, simply knowing that a variant exists (regardless of exact zygosity) is possible with high confidence. However, applications requiring precise dosage information (e.g., distinguishing 1/4 from 2/4 in tetraploids) may require higher coverage or more specialized algorithms. Although genotyping accuracy still has room for improvement, the ability to generate haplotype-aware assemblies suggests that phasing-oriented strategies, such as local de novo assembly, would be particularly effective. Nonetheless, the ability to get >95% recall and precision for SNV discovery in a tetraploid like potato using ∼30× long-read data is a significant step forward for plant genomics.

A key limitation we observed is that genome complexity, especially repetitive content and structural variation, continues to hinder accurate variant calling, even with high-fidelity long reads. In *F. vesca*, *S. tuberosum*, and *Z. mays*, false positives frequently arose from reference bias and misalignments in repeat-rich regions. This indicates that read length alone is not sufficient, and further improvements in alignment and reference models are needed. Mapping quality (MQ) scores generally reflected genome complexity across species, with higher values in *F. vesca* and lower in *Z. mays* (Table S2). This suggests that aligners adjust for repeat content when scoring reads. However, many false-positive variants in maize were still supported by reads with high MQ, implying that local repeat-induced ambiguity is not fully captured. Reads from sample-specific or absent regions may misalign to homologous repeats elsewhere, generating confident but incorrect placements.

These insights have practical implications for crop genomics and breeding. A single linear reference is a bottleneck: if an accession’s genome differs significantly (as in maize), the aligner contorts reads to fit the reference, leading to mistakes. Graph genome representations, which incorporate alternate haplotypes, can in principle reduce reference bias by allowing reads to align to the correct version of a sequence present in the population [27, 28]. Even with a graph-based pangenome, it remains critical that the pangenome includes sequences sufficiently close to the target genome. Otherwise, misalignment issues may still arise, leading to genotyping errors or false-positive variant detection, despite the use of a pangenome framework.

It is also important to note that reference bias in long-read data may differ in nature from that reported in short-read studies. Previous analyses, such as those based on RNA-seq in maize [29], have primarily highlighted reduced mapping recall when reads originate from haplotypes divergent from the reference. In contrast, our results indicate that with long reads, the primary concern is not loss of recall but a reduction in precision, driven by confident yet incorrect alignments to homologous regions. This shift in error mode underscores the need for bias-aware strategies tailored to the properties of long-read sequencing.

From an applied perspective, our findings have practical ramifications for crop genomics and breeding. In maize and other large-genome cereals, reliance on a single reference genome can introduce erroneous variant calls due to substantial structural divergence. The development of pangenome references, already underway in numerous crop research communities, represents a promising path forward. Our results provide quantitative support for this direction and highlight the degree of improvement required. A pangenome approach will likely be the way forward; our data provides quantitative evidence of how much improvement is needed. In essence, the accuracy of small variant detection in complex plant genomes is now largely a function of reference quality and completeness. As more reference genomes become available, variant calling accuracy is expected to improve correspondingly.

A critical consideration is that the use of a pangenome reference does not automatically eliminate the problems observed in this study. If the sample genome contains sequences that are absent from the pangenome graph, the aligner may still assign reads to graph nodes with only partial or local similarity. This can perpetuate the same alignment errors and reference bias that the pangenome is intended to mitigate. Therefore, the pangenome must capture a sufficiently wide range of genomic diversity. Equally important, mapping and variant-calling algorithms must incorporate mechanisms to detect and correct misalignments and false-positive calls, even in the presence of a comprehensive pangenome reference.

## Conclusions

In this study, we systematically benchmarked variant callers for complex genomes such as polyploid and highly repetitive genomes using high-accuracy long reads across four species. Our evaluation revealed that while existing tools perform well for simple diploid genomes, their accuracy significantly deteriorates in repetitive contexts, particularly in *Zea mays*. We also found that commonly used quality filters, including mapping quality (MQ), may fail to eliminate false positives, depending on genome complexity.

Our findings emphasize that addressing reference bias during the alignment step is critical, and both variant caller developers and users should carefully consider its impact in polyploid analyses. Future work should focus on designing polyploid-aware algorithms that integrate alignment uncertainty and ploidy-aware genotyping for improved variant discovery.

## Methods

### Data sets and genome references

#### Human polyploid benchmarking data

We used the well-characterized NA12878 Ashkenazi trio from the Genome in a Bottle (GIAB) project for our human variant calling benchmarks. High-quality variant truth sets are available for the child (HG002) and parents (HG003, HG004) on the GRCh38 reference genome. We obtained PacBio HiFi whole-genome sequencing reads (CCS reads) for these samples from public human pangenome consorthium releases (coverage ∼30+× per genome) [2, 30]. The GRCh38 reference assembly (with decoy sequences and ALT contigs removed) was used for alignment and evaluation. High-confidence variant calls for HG002, HG003, HG004 (GIAB v4.2.1) were downloaded to serve as ground truth [31].

#### Plant genomes and sample selection

We selected three plant species to represent a gradient of genome complexity. *F. vesca* (woodland strawberry) is a diploid plant with a small genome (∼240 Mb, 7 chromosomes); we used the *F. vesca* reference genome ver. 6 [32] and used a recently released independent assembly as a proxy for the sample under investigation. *S. tuberosum* (potato) is an autotetraploid (4n=4x=48) with a ∼840 Mb genome; we used the DM1-3 516 R44 v6.1 reference (a doubled monoploid line [33]) and tetraploid variety Otava that has been assembled in a haplotype aware methodology [21]. *Z. mays* (maize) is diploid (2n=20) with an ∼2.3 Gb genome rich in repeats (∼88% transposable elements); we used the B73 ver. 5 reference genome and selected another inbred line Mo17 as a target genome that has been assembled at the telomere-to-telomere scale [20].

#### Construction of synthetic polyploids and truth sets

For the human benchmark, we created synthetic polyploid truth sets by leveraging the known high-confidence variants of the GIAB trio. The tetraploid truth set was defined by taking the union of variant sites from HG003 and HG004 (the two parents), restricted to regions where both had confident calls. Similarly, the hexaploid truth set was the union of HG002, HG003, HG004 variant sites. This approach follows Cooke et al. (2022) who merged GIAB callsets to benchmark polyploid calling [10]. Because the GIAB truth data are highly reliable, we treated this union as ground truth for variant existence. We simulated the sequencing of these polyploids by combining reads: all HiFi reads from HG003 and HG004 were merged into one dataset for the tetraploid, and adding HG002 reads to that for the hexaploid. The read depth per haplotype was thereby roughly equal (e.g., ∼15× per haplotype in the tetraploid, summing to 60× total coverage).

#### Construction of plant benchmark sets

For the plant genomes, establishing a truth set is more challenging due to lack of a priori ground truth. We aligned the high-quality assemblies to the reference genome and defined small variants located within the most confidently aligned regions as benchmarks for downstream analyses [17]. We used dipcall ver. 0.3 for the analysis. Because dipcall was originally developed for diploid genomes, we extended its functionality to enable similar analyses for the tetraploid potato genome. *F. vesca* is self-compatible and known for its low heterozygosity. Additionally, a haplotype-resolved genome assembly was recently released by the Tree of Life Project. We utilized this haplotype-resolved assembly for comparative analysis, using the representative accession, Hawaii-4, as the reference genome [34]. For *S. tuberosum*, we performed a similar analysis by aligning the recently reported haplotype-resolved assembly of the Otava cultivar [21] to the reference genome of doubled monoploid DM1-3 516 R44 [33]. For *Z. mays*, we took advantage of the availability of a reference-quality telomere-to-telomere assembly of the Mo17 inbred [20]. We aligned the Mo17 assembly to the 5^th^ version of B73 reference to derive a comprehensive set of variants. Mo17 is expected to exhibit very low heterozygosity, so variants were treated as homozygous in our analyses. Although a small fraction of heterozygous sites may remain [35], their frequency is assumed to be negligible within the accuracy range assessed in this study. For *F. vesca* and *Z. mays*, we adopted the minimap2-x asm5 preset, as applied by previous studies [36]. For *S. tuberosum*, due to its higher sequence divergence [21], the alignment stringency was relaxed by using the-x asm20 preset. All other parameters followed the default settings of dipcall. For diploid species (*F. vesca* and *Z. mays*), we used Merfin to assess variant based on *k*-mer multiplicity independently [37]. Merfin was not applied for potato due to its complex polyploid *k*-mer context. In all cases, we restricted our precision/recall calculations to regions deemed high confidence for truth. These evaluation masks ensure that we are not unfairly penalizing the variant callers for variants that are essentially unresolvable or not represented in our truth data. Because both the truth assembly and the benchmarking reads derive from the same individual, sample-specific insertions are identical to the assembly sequence; therefore, no SNP or indel variants exist within non-alignable regions, and their exclusion does not affect the reported metrics.

#### HiFi simulation for the *Zea mays* genome

To simulate HiFi from ccs calling step, subreads were simulated from the *Z. mays* inbred line Mo17 to a depth of 30 × using PBSim3 [38] using quality score model under whole-genome-shotgun settings (mean template length = 12,000 bp; pass-num = 10). Circular consensus sequence (CCS) reads were then generated from the simulated subreads with ccs command in SMRT Link 12.

### Alignment and variant calling pipeline

#### Read alignment

We aligned long reads to their respective reference genomes using minimap2 ver. 2.24 [39]. For PacBio HiFi reads, we used the same parameters applied in pbmm2, which employs sensitive settings for high-accuracy reads. Alignments were output in SAM/BAM format and sorted and indexed with samtools. Distribution of mapping quality was computed using stats subcommand in samtools ver. 1.17.

#### Variant calling

We evaluated two variant calling tools that support polyploid genotyping: GATK HaplotypeCaller (from GATK ver. 4.1.4) and FreeBayes (v1.3.6) [40, 41]. GATK HaplotypeCaller was executed with ploidy set to the appropriate value for each sample (i.e.,-ploidy 2 for diploids) with recommended filtering parameters [2]. FreeBayes was run with “-=“ and the--ploidy parameter similarly set (FreeBayes documentation states that hard input cut-offs seldom improve accuracy and recommends post-hoc site filtering, see below), and the other parameters were set to default. For the human trio-derived datasets, we ran variant calling in three scenarios: diploid (ploidy=2), tetraploid (ploidy=4), and hexaploid (ploidy=6) on the synthetic merged read sets. We did not include DeepVariant [42] in our pipeline because it is designed for diploid germline and cannot output polyploid genotypes. For reference, we did run DeepVariant on the tetraploid *S. tuberosum* datasets as a supplementary experiment, and it showed some notable bias in tetraploid case (Table S3 and S4). Therefore, we focus on GATK/FreeBayes for consistency across ploidies. Octopus (v0.7.4) was evaluated using the default short-read mode on PacBio HiFi data, as no officially supported long-read configuration was available. A separate test using the developmental configuration for PacBio (PacBioCCS.config) was also attempted but did not produce stable output due to repeated runtime errors. Under short-read mode, the tool executed stably but exhibited suboptimal variant calling performance (recall 1.3%, precision 52.4%, F1 score 2.6% for *F. vesca*). Given the lack of robust support for long-read data and the suboptimal performance, Octopus was excluded from final benchmarking comparisons.

To ensure fair runtime and memory comparisons, we benchmarked all tools on ∼10% of the genome (uniformly distributed) using a single workstation (Intel Xeon CPU, 256 GB RAM, 10 threads) to avoid variability introduced by heterogeneous computing environments. Full-genome benchmarking was not conducted due to significant computational time requirements. The results are intended to provide representative estimates (Table S6).

#### Variant filtering

The raw variant calls were filtered to produce high-confidence call sets for evaluation. We applied hard filters on variant quality metrics. For GATK, we applied the community-recommended per-type QD hard filters: variants were filtered if QD < 2 for SNVs and indels greater than 1 bp, and QD < 5 for 1-bp indels. For FreeBayes, pilot tests using chr1 of HG002 confirmed that raising the MQM threshold to 60 or applying per-type QD filters, as done in GATK, consistently resulted in slightly reduced F1 scores (Table S8). We therefore applied the same hard-read filters for FreeBayes as previously described [10]. We also did not evaluate any variant calls in regions of low reliability: specifically, we extracted variants with intervals that had reliable alignments between reference and target genome assemblies using minimap2 called by dipcall.

#### Evaluation metrics and analysis

We calculated precision, recall, and F1-score for variant calls by comparing the caller’s VCF to the truth set VCF for each sample. RTG Tools vcfeval was applied for benchmarking. Genotype concordance (the correctness of allele dosage) was evaluated for the polyploid human samples by comparing the called genotype to the truth genotype. To investigate error causes, we manually inspected alignments and variant calls in IGV for a number of discordant sites, especially clusters of false positives in the plant genomes.

By combining controlled experiments on human data with real-world tests on plants, our methods provide a general template for assessing variant calling in non-model genomes, and our pipeline can be readily adapted to other species or sequencing technologies in future work.

## Data availability

All analysis code, including custom scripts for merging truth sets and evaluating polyploid calls, is available in a public GitHub repository. Code for evaluating polyploid calls is available at github (https://github.com/yfukasawa/dipcall-poly). Code, snakemake pipeline, and docker image to run them for reproducing the results are available at https://github.com/yfukasawa/long-read-wgs-polyploid-benchmark.

## Funding

This study was partly supported by JSPS KAKENHI Grant Number 23K19338 to YF and the Utsunomiya University Strawberry Project.

## Acknowledgments

Computations were partially performed on the NIG supercomputer at ROIS National Institute of Genetics.

## Contributions

YF designed the research, performed experiments, conducted analyses, interpreted results, and wrote the manuscripts.

## Ethics declarations

### Ethics approval and consent to participate

Not applicable.

### Consent for publication

All authors have read and agreed with the publication.

### Competing interests

The authors declare no competing interests.

## Notes

### Competing Interest Statement

The authors have declared no competing interest.

### Summary of Updates

In this version, we added our benchmarking results regarding computational time using improved pipelines. We also revised the discussion on mapping quality filters, clarified the impact of reference bias, and added the conclusion for clarity. We also revised the title to better reflect the scope and generality of the study.

