## Supplemental materials for "Benchmarking long-read variant calling in diploid and polyploid genomes: insights from human and plants"

### Supplementary Figures

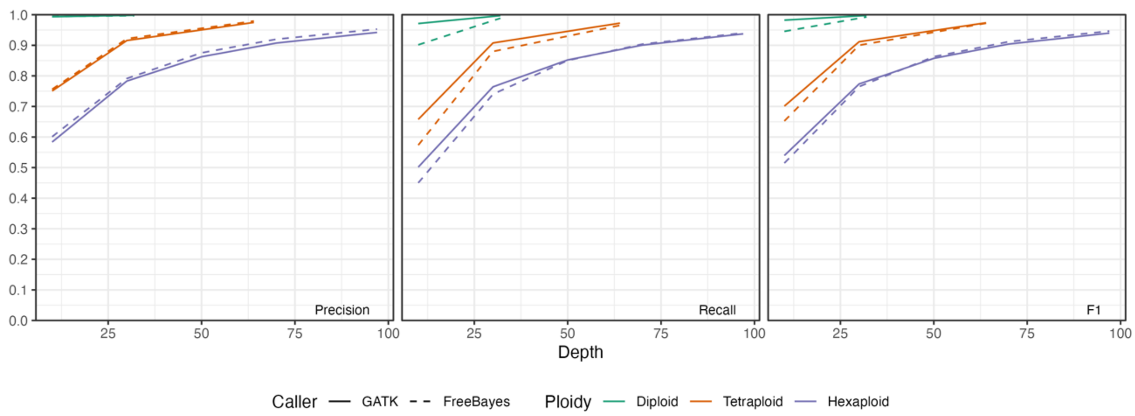


Figure S1 Performance of SNV genotyping on the synthetic human polyploid genome using high-accuracy long reads. Precision, recall, and F1 scores are presented.

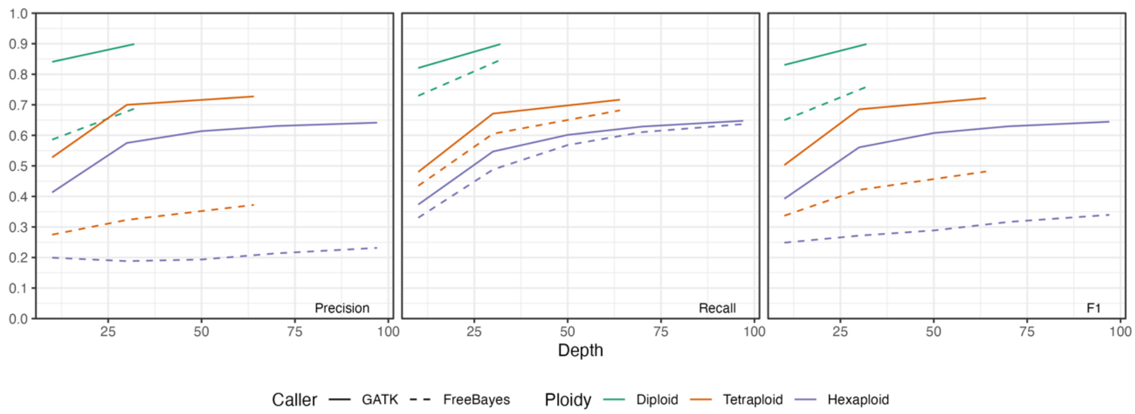


Figure S2 Performance of indel genotyping on the synthetic human polyploid genome using high-accuracy long reads. Precision, recall, and F1 scores are presented.

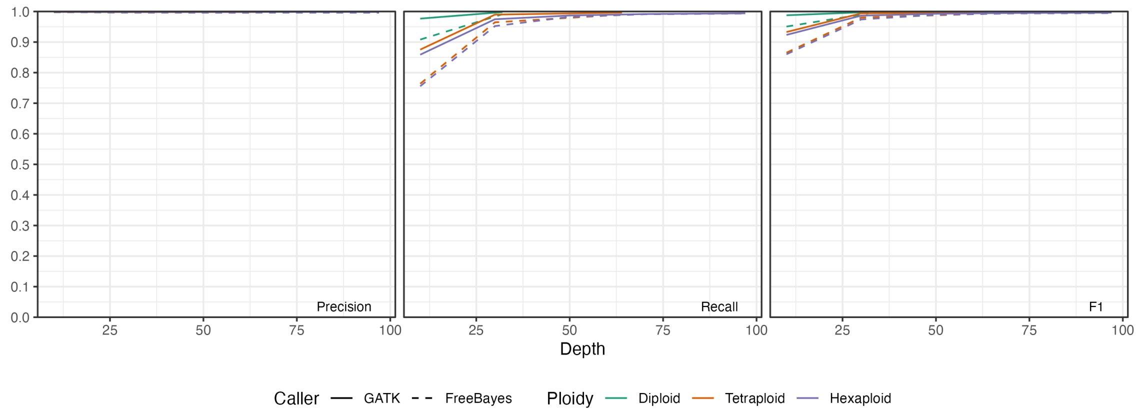


Figure S3 Performance of SNV detection on the synthetic human polyploid genome using high-accuracy long reads. Precision, recall, and F1 scores are presented.

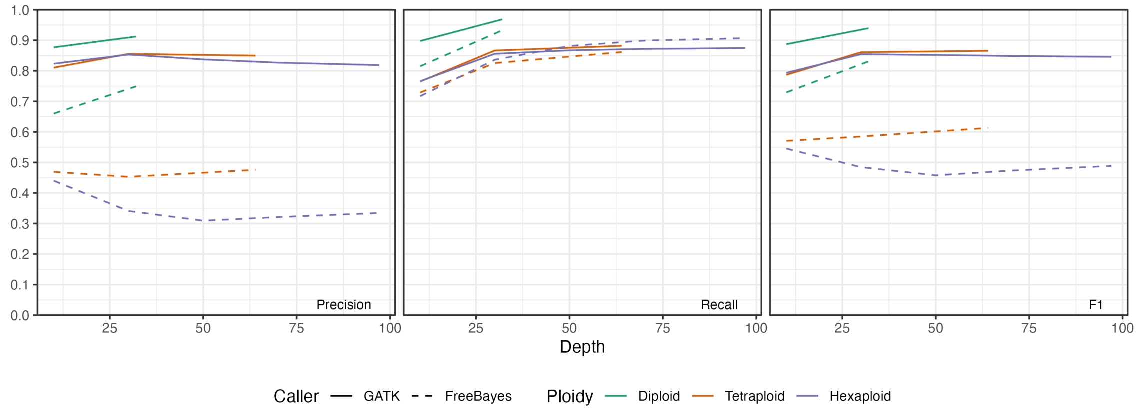


Figure S4 Performance of indel detection on the synthetic human polyploid genome using high-accuracy long reads. Precision, recall, and F1 scores are presented.

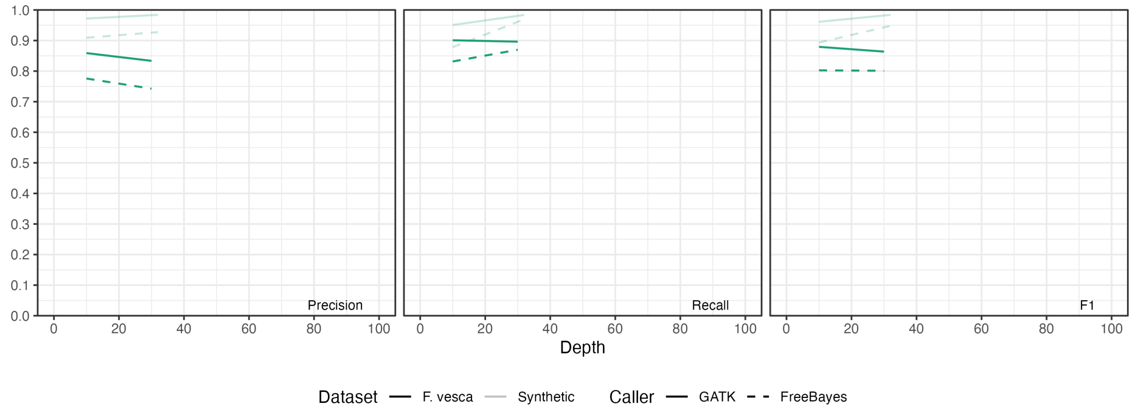

Figure S5 Performance of small variant genotyping on *F. vesca* genome using high-accuracy long reads. Precision, recall, and F1 scores are presented.

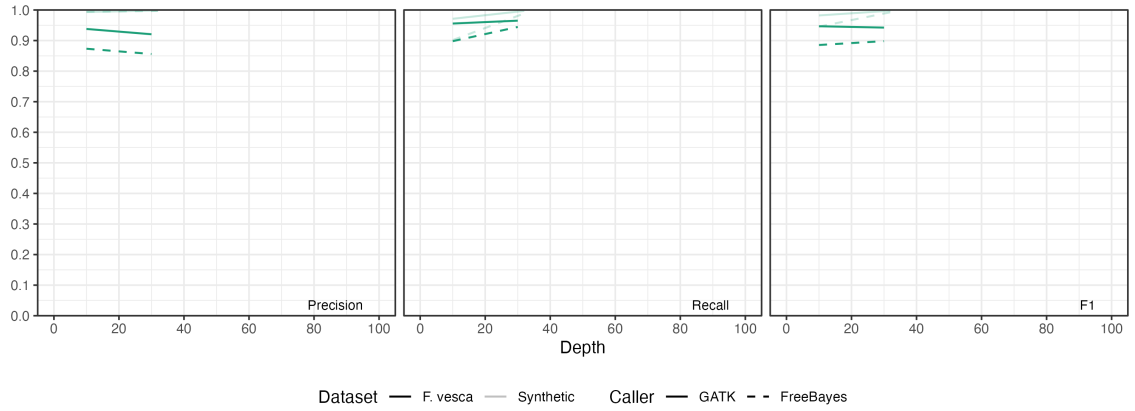


Figure S6 Performance of SNV genotyping on *F. vesca* genome using high-accuracy long reads. Precision, recall, and F1 scores are presented.

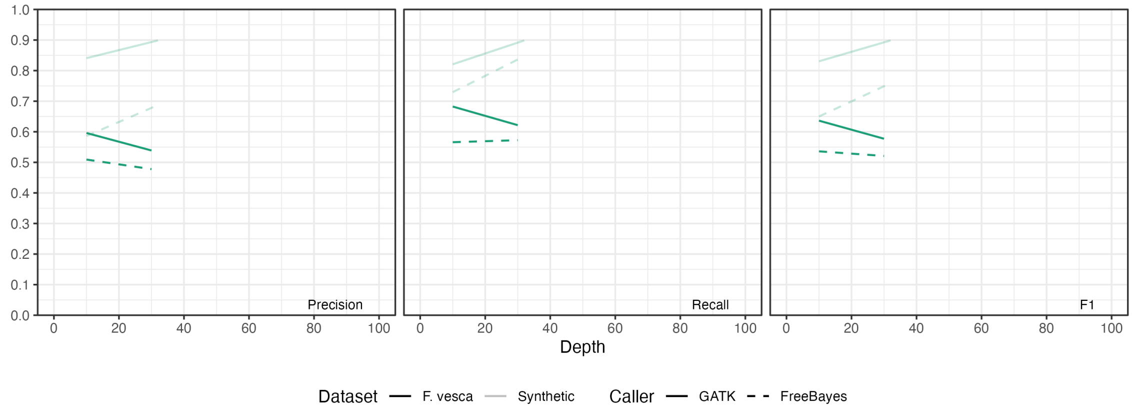


Figure S7 Performance of indel genotyping on *F. vesca* genome using high-accuracy long reads. Precision, recall, and F1 scores are presented.

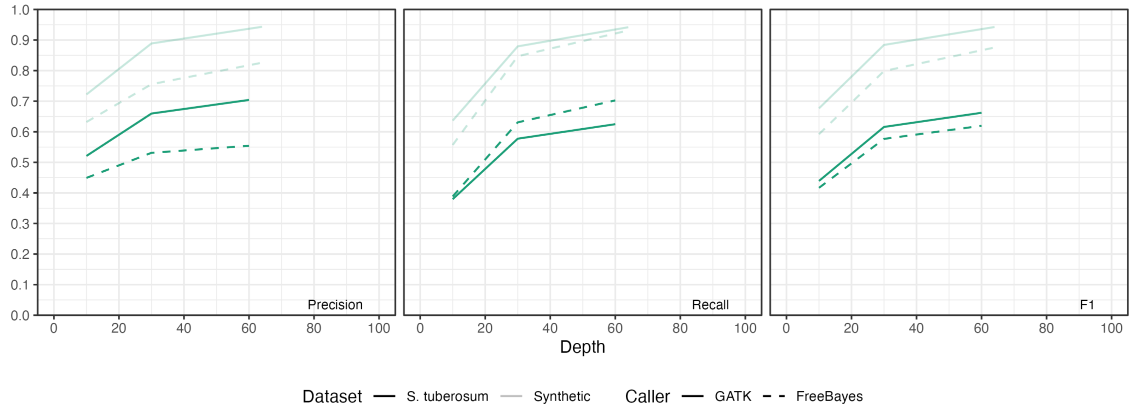

Figure S8 Performance of small variant genotyping on *S. tuberosum* genome using high-accuracy long reads. Precision, recall, and F1 scores are presented.

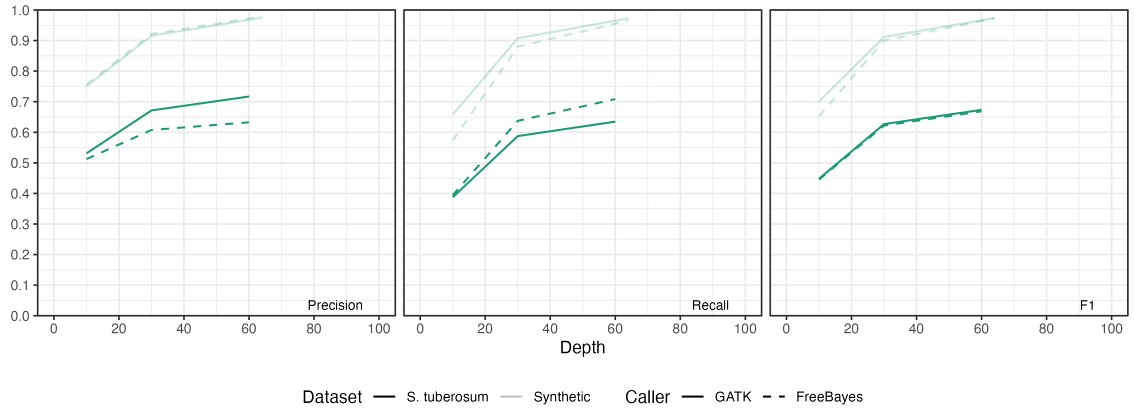


Figure S9 Performance of SNV genotyping on *S. tuberosum* genome using high-accuracy long reads. Precision, recall, and F1 scores are presented.

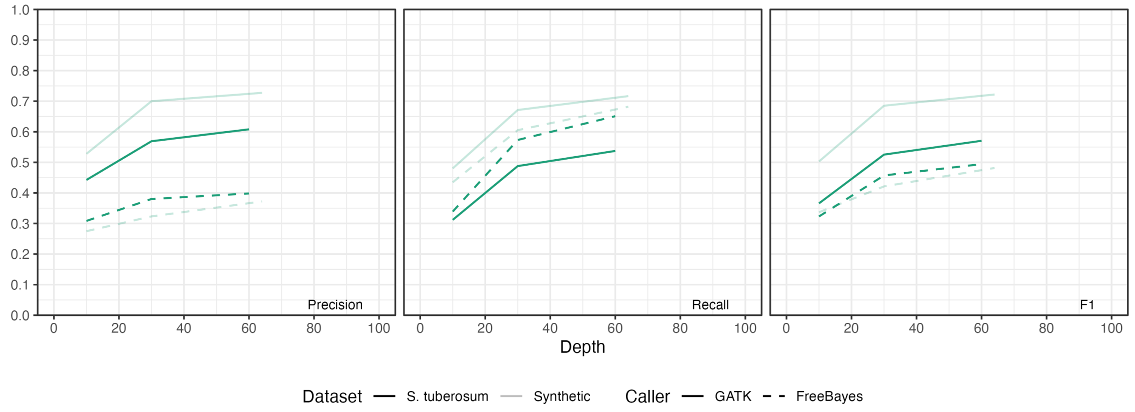


Figure S10 Performance of indel genotyping on *S. tuberosum* genome using high-accuracy long reads. Precision, recall, and F1 scores are presented.

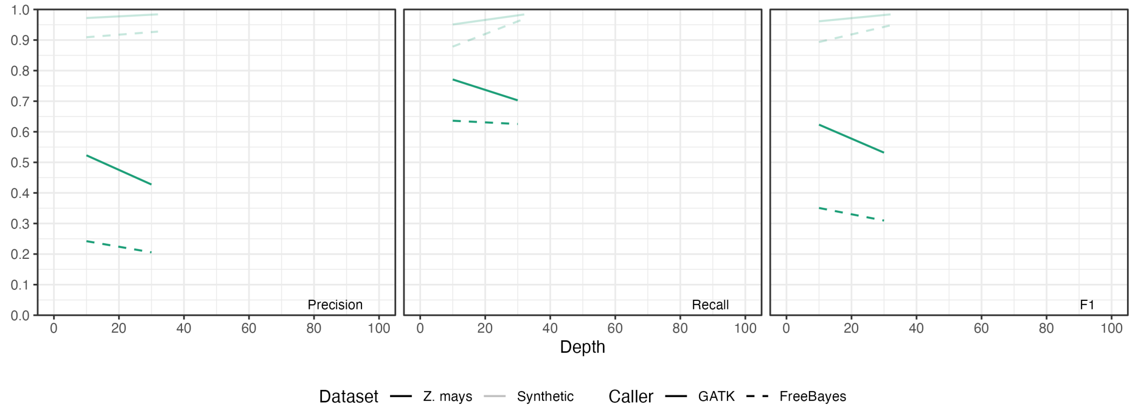

Figure S11 Performance of small variant genotyping on *Z.mays* genome using high-accuracy long reads. Precision, recall, and F1 scores are presented.

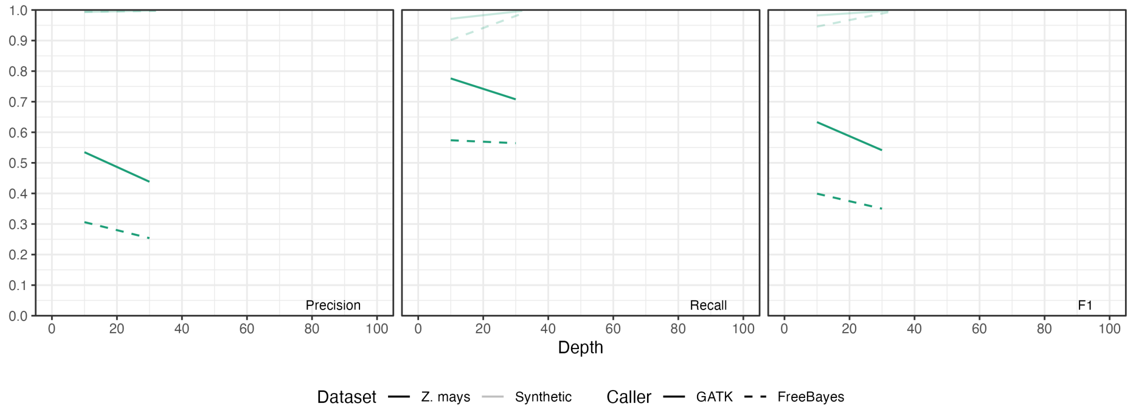


Figure S12 Performance of SNV genotyping on *Z.mays* genome using high-accuracy long reads. Precision, recall, and F1 scores are presented.

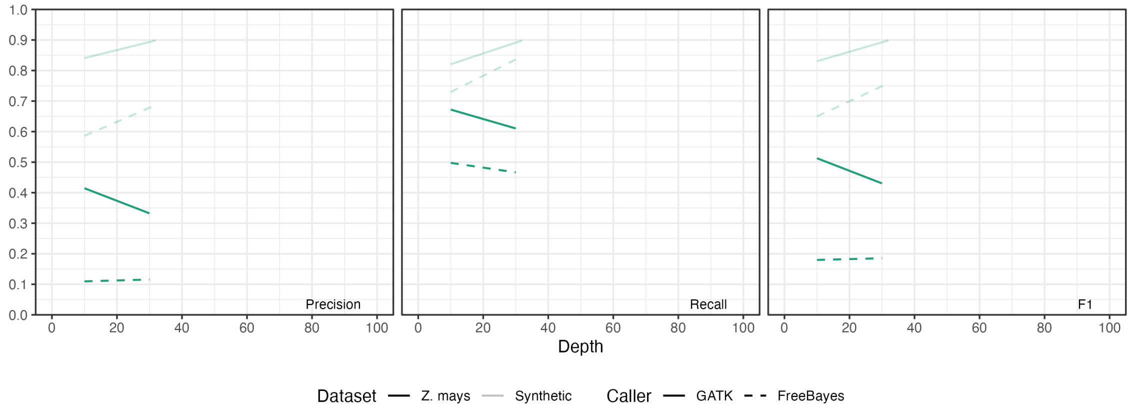


Figure S13 Performance of Indel genotyping on *Z.mays* genome using high-accuracy long reads. Precision, recall, and F1 scores are presented.

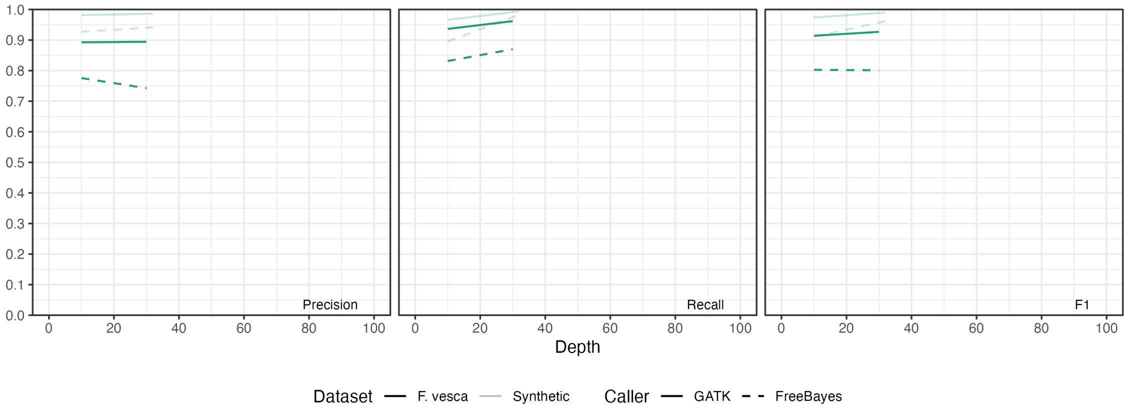


Figure S14 Performance of small variant detection on the *F. vesca* genome using high-accuracy long reads.

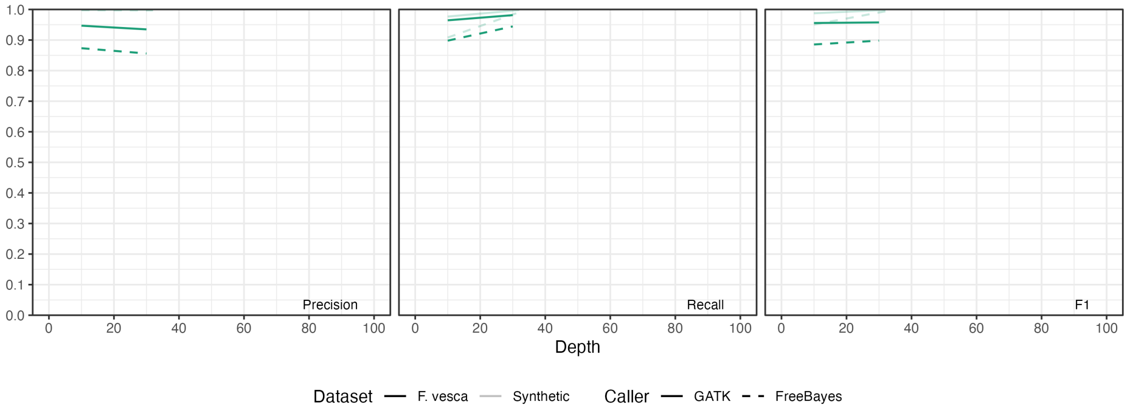


Figure S15 Performance of SNV detection on the *F. vesca* genome using high-accuracy long reads.

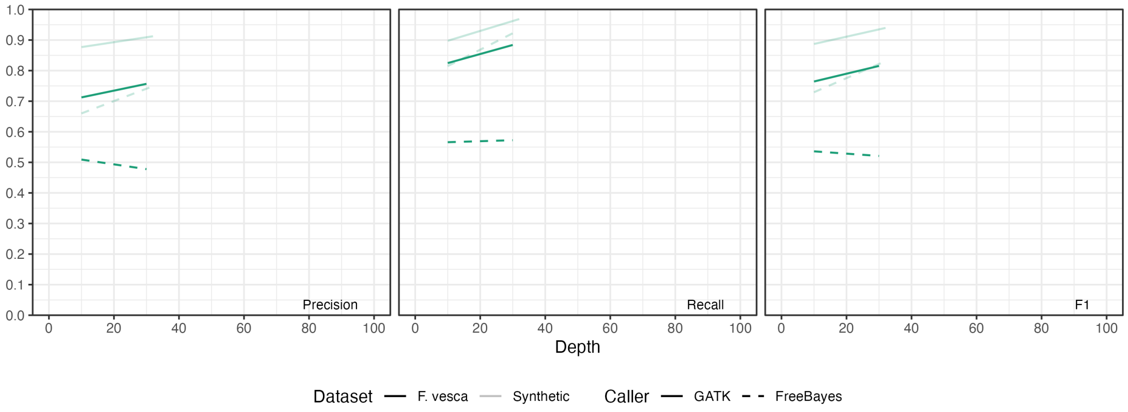

Figure S16 Performance of Indel detection on the *F. vesca* genome using high-accuracy long reads.

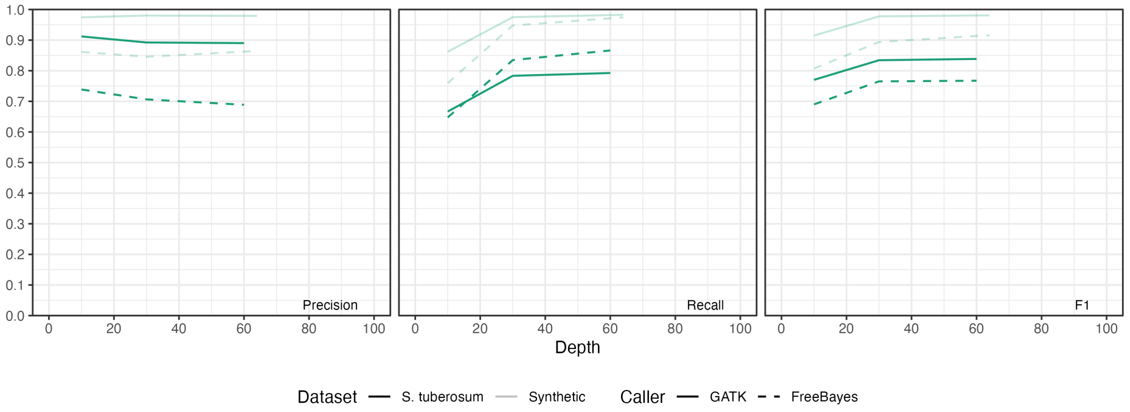


Figure S17 Performance of small variant detection on the *S. tuberosum* genome using high-accuracy long reads.

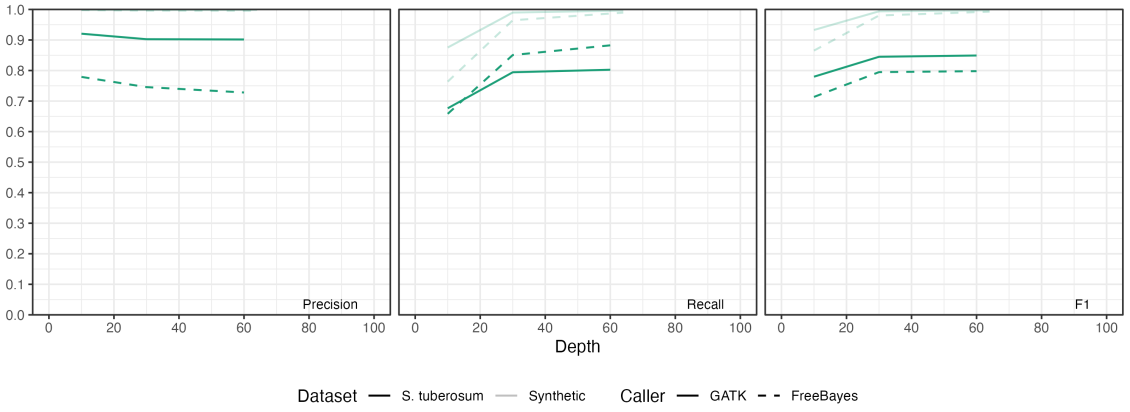


Figure S18 Performance of SNV detection on the *S. tuberosum* genome using high-accuracy long reads.

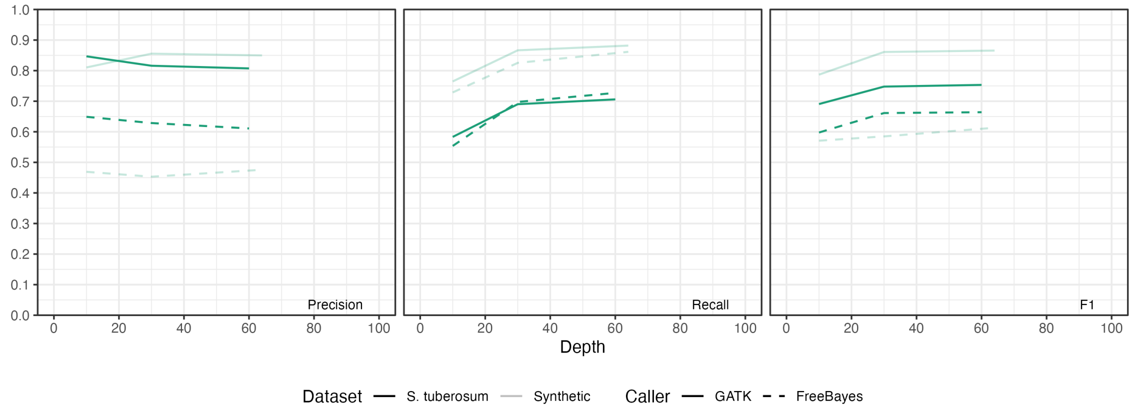


Figure S19 Performance of Indel detection on the *S. tuberosum* genome using high-accuracy long reads.

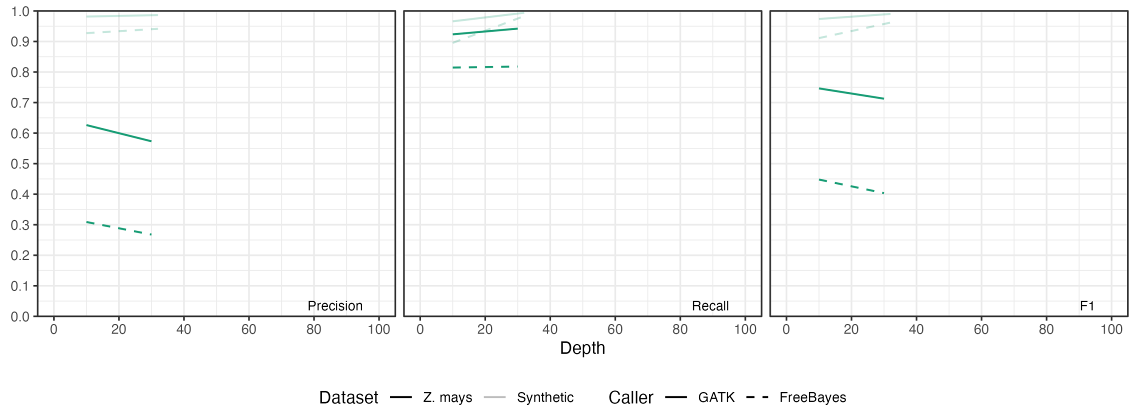


Figure S20 Performance of small variant detection on the *Z. mays* genome using high-accuracy long reads.

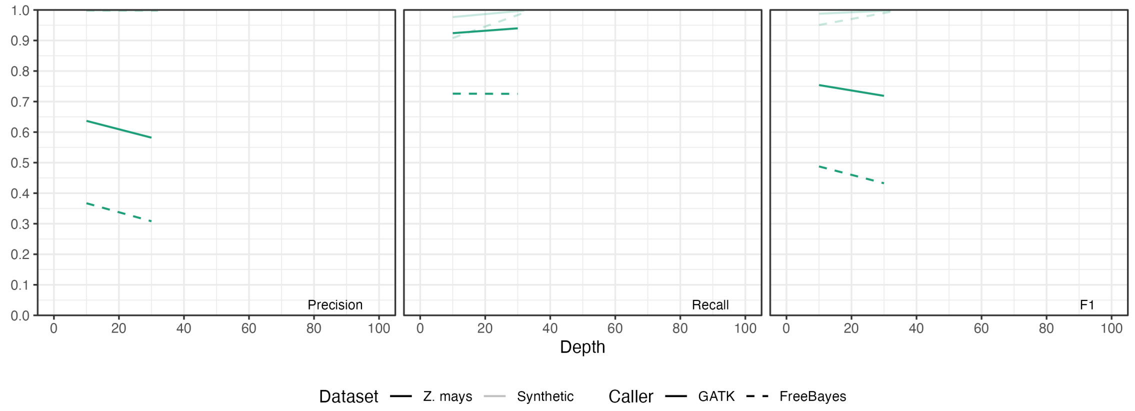


Figure S21 Performance of SNV detection on the *Z. mays* genome using high-accuracy long reads.

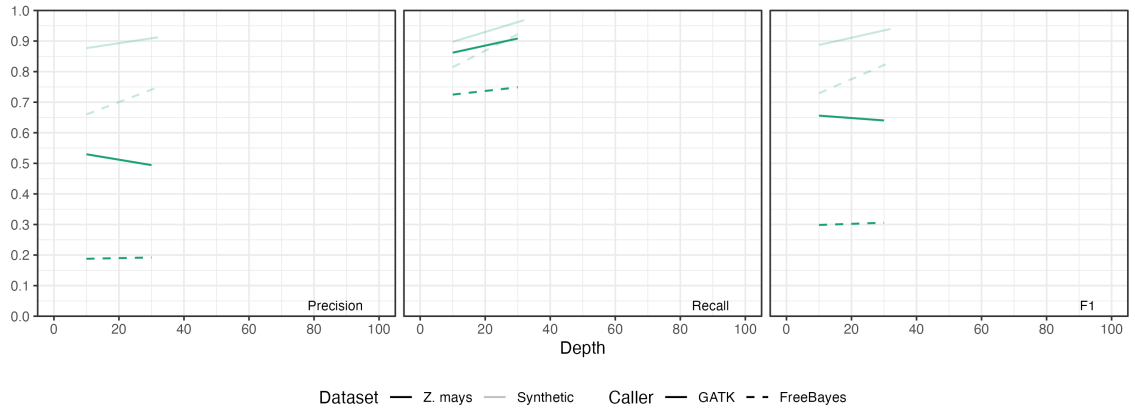


Figure S22 Performance of Indel detection on the *Z. mays* genome using high-accuracy long reads.

### Supplementary Tables

Table S1. List of accession numbers or link for the PacBio HiFi sequencing datasets analyzed in this study.

(Accession details are provided in a separate Excel file available as supplementary material.)

Table S2. Mapping statistics for the datasets used in this study, including Homo sapiens (HG002), Fragaria vesca (drFraVesc1), Solanum tuberosum (Otava), and Zea mays (Mo17). The BAM file with the highest sequencing depth for each respective sample was selected for quality validation.

(Detailed mapping statistics are provided in a separate Excel file available as supplementary material.)

Table S3. Comprehensive performance metrics for small-variant detection across all samples and variant callers evaluated in this study. Each row corresponds to a species, and each column reports evaluation metrics including precision, recall, and F1 score for all, SNVs and indels. Results are shown for each variant caller and ploidy configuration where applicable. Benchmarking was performed using the corresponding truth set for each genome. All performance values reflect post-filtered variants within confident regions. See Methods for benchmarking criteria.

(Detailed statistics are provided in a separate Excel file available as supplementary material.)

Table S4. *k*-mer-based validation of detected variants in diploids using Merfin

| | Validated | | | Total | | |
|---|---|---|---|---|---|---|
| | SNP | Indel | Other | SNP | Indel | Other |
| *F. vesca* | 233,607 | 55,828 | 0 | 233,913 | 56,210 | 0 |
| Validated % | 99.90% | 99.30% | | | | |
| *Z. mays* | 2,575,758 | 245,962 | 1 | 2,589,324 | 248,328 | 1 |
| Validated % | 99.50% | 99.10% | | | | |

Table S5. Performance of small variant detection by GATK and DeepVariant on the *S. tuberosum* genome using high-accuracy long reads

| **GATK** | depth | Precision | Sensitivity | F-measure |
|---|---|---|---|---|
| *S. tuberosum* | 10x | **0.912** | 0.6666 | 0.7702 |
| *S. tuberosum* | 30x | 0.8923 | 0.7834 | 0.8343 |
| *S. tuberosum* | 60x | 0.8902 | **0.7925** | **0.8385** |
| **DeepVariant** | | | | |
| *S. tuberosum* | 10x | 0.9492 | 0.6485 | 0.7706 |
| *S. tuberosum* | 30x | 0.9572 | **0.7167** | **0.8197** |
| *S. tuberosum* | 60x | **0.9582** | 0.6576 | 0.7799 |

Table S6. Number of false negatives in variant detection by GATK and DeepVariant on the *S. tuberosum* genome, stratified by genotype at 60× sequencing depth

| Genotype | #FNs (DeepVariant) | #FNs (GATK) | The total number |
|---|---|---|---|
| AAAa | 1078242 (59.3%) | 654961 (36.0%) | 1818863 |
| AAaa | 275882 (25.7%) | 197695 (18.5%) | 1071719 |
| Aaaa | 151736 (15.9%) | 82340 (8.6%) | 952523 |
| aaaa | 8579 (1.5%) | 19636 (3.4%) | 579396 |

Table S7. Computational time and memory usage for each caller. Benchmark was conducted on ~10% of the genome using a single workstation. Max RSS was measured per process, and the peak value among all concurrent processes was reported to represent each caller's memory demand.

| Caller | Species | Ploidy | Depth | Wall time (s) | CPU time (s) | Max RSS (MB) |
|---|---|---|---|---|---|---|
| GATK | *Fragaria vesca* | *2* | 10x | 314.32 | 400.29 | 5061.2 |
| GATK | *Fragaria vesca* | *2* | 30x | 547.34 | 644.21 | 4018.3 |
| GATK | *Zea mays* | *2* | 10x | 7145.95 | 8197.62 | 4232.7 |
| GATK | *Zea mays* | *2* | 30x | 13719.83 | 15151.91 | 3875.8 |
| GATK | *Solanum tuberosum* | *4* | 10x | 8295.91 | 8615.93 | 4987.5 |
| GATK | *Solanum tuberosum* | *4* | 30x | 12150.32 | 12584.62 | 4286.9 |
| GATK | *Solanum tuberosum* | *4* | 60x | 15333.96 | 15998.02 | 3408.6 |
| GATK | *Homo sapiens* | *6* | 10x | 10048.31 | 4484.03 | 4256.8 |
| GATK | *Homo sapiens* | *6* | 30x | 21717.4 | 8473.08 | 3369.4 |
| GATK | *Homo sapiens* | *6* | 60x | 31836.92 | 15275.62 | 3195.1 |
| GATK | *Homo sapiens* | *6* | 70x | 37392.96 | 25266.48 | 3267.6 |
| GATK | *Homo sapiens* | *6* | 90x | 58383.12 | 51860.75 | 5643.6 |
| FreeBayes | *Fragaria vesca* | *2* | 10x | 243.49 | 225.32 | 366.4 |
| FreeBayes | *Fragaria vesca* | *2* | 30x | 800.47 | 795.62 | 972.6 |
| FreeBayes | *Zea mays* | *2* | 10x | 18569.91 | 18461.82 | 863.5 |
| FreeBayes | *Zea mays* | *2* | 30x | 58900.31 | 58674.15 | 806 |
| FreeBayes | *Solanum tuberosum* | *4* | 10x | 6022.81 | 5853.31 | 187.9 |
| FreeBayes | *Solanum tuberosum* | *4* | 30x | 17282.8 | 17256.43 | 690.6 |
| FreeBayes | *Solanum tuberosum* | *4* | 60x | 29278.88 | 29254.79 | 1190.5 |
| FreeBayes | *Homo sapiens* | *6* | 10x | 2990.81 | 1923.94 | 688.8 |
| FreeBayes | *Homo sapiens* | *6* | 30x | 8113.17 | 7324.45 | 564.3 |
| FreeBayes | *Homo sapiens* | *6* | 60x | 9371.91 | 8656.49 | 584.8 |
| FreeBayes | *Homo sapiens* | *6* | 70x | 12924.51 | 12565.68 | 490.4 |
| FreeBayes | *Homo sapiens* | *6* | 90x | 13976.63 | 13571.25 | 1433.1 |

Table S8. Assessment of the impact of MQM and QD on variant calling performance in FreeBayes using HG002 chromosome 1

| MQM | Depth | Category | Precision | Sensitivity | F-measure |
|---|---|---|---|---|---|
| 1 | 10 | MQM1-Depth10 | 0.9013 | 0.8776 | 0.8893 |
| 1 | 32 | MQM1-Depth32 | 0.9202 | **0.9665** | **0.9428** |
| 60 | 10 | MQM60-Depth10 | 0.9025 | 0.874 | 0.888 |

| | | | | | |
|---|---|---|---|---|---|
| 60 | 32 | MQM60-Depth32 | 0.9227 | 0.9586 | 0.9403 |
| 1 | 10 | MQM1-Depth10 with GATK-like filter | 0.936 | 0.8454 | 0.8884 |
| 1 | 32 | MQM1-Depth32 with GATK-like filter | 0.9383 | 0.906 | 0.9218 |
| 60 | 10 | MQM60-Depth10 with GATK-like filter | 0.9369 | 0.842 | 0.8869 |
| 60 | 32 | MQM60-Depth32 with GATK-like filter | **0.9401** | 0.8987 | 0.9189 |
